# Substrate adhesion determines migration during mesenchymal cell condensation in chondrogenesis

**DOI:** 10.1101/2022.05.17.492260

**Authors:** Ignasi Casanellas, Hongkai Jiang, Carolyn M. David, Yolanda Vida, Ezequiel Pérez-Inestrosa, Josep Samitier, Anna Lagunas

## Abstract

Effective cartilage development relies on the successful formation of mesenchymal cell condensates. Mesenchymal condensation is a prevalent morphogenetic transition, which involves the upregulation of the adhesive extracellular glycoprotein fibronectin (FN). During condensation, there is an active directional migration of cells from the surrounding loose mesenchyme towards regions of increasing matrix adherence (the condensation centers). In this study, we live imaged the first 40 h of mesenchymal condensation during chondrogenesis on nanopatterns of the cell-adhesive peptide arginine-glycine-aspartic acid (RGD), present in FN. Results show cell-substrate adhesions modulate both single-cell and collective cell migration during mesenchymal condensation. Single cell tracking analysis showed that substrate adhesion determines the migration mode, protrusion formation and the directionality of the cell movement. Cells on the more adhesive nanopatterns presented traits among amoeboid and mesenchymal modes of migration facilitating a more directional movement and reducing contact inhibition of locomotion (CIL), which allows merging and condensation. Inhibition experiments demonstrated that neural cadherin (*N*-Cad) is required in cell-cell interactions, enabling cells to coordinate their movement and directionality in a multicellular environment and to maintain the group cohesiveness during migration. Altogether, this contributes to create a sufficiently dynamic scenario, in which there is a balance between cell-substrate and cell-cell adhesions for condensates to grow. Our results provide a framework for the regulation of single and collective cell migration during mesenchymal condensation, through nanoscale cell-substrate adherence.

**Summary statement:** The fine tuning of substrate adherence through nanopatterning allows control of mesenchymal cell migration and determines condensation during chondrogenesis *in vitro*.

## 1. Introduction

Condensations of mesenchymal cells are present during development of many tissues and organs, including cartilage and bone. In osteochondral development, condensation is a transient phase in which the size and position of each cartilage anlage is defined (DeLise et al., 2000). An abnormal condensation process results in developmental defects (Atchley and Hall, 1991; Giffin et al., 2019). Several inductive epithelial signaling molecules have been identified as initiators of mesenchymal condensation (Giffin et al., 2019). Transforming growth factor (TGF-β) induces the condensation of mesenchymal cells at high density through the upregulation of the adhesive extracellular glycoprotein fibronectin (FN) (Miura and Shiota, 2000; Singh and Schwarzbauer, 2012). It has been proposed that condensation results from extracellular matrix (ECM)-driven translocation and directional migration of cells from the surrounding loose mesenchyme to the condensed region (Newman and Tomasek, 1996). During condensation, active mesenchymal cell migration towards condensation centers (regions of increased cell–FN adhesive interactions) causes an increase in mesenchymal cell-packing density without cell proliferation (Janners and Searls, 1970), in which cells acquire epithelioid properties by the upregulation of cell-cell adhesion molecules (Oberlender and Tuan, 1994). Cell gathering favors the establishment of cell-cell contacts through the cell adhesion molecules *N*-Cad, neural adhesion molecule (N-CM) (Widelitz et al., 1993), and gap junctions (GJs) that facilitate cell coordination within de cohort (Coelho and Kosher, 1991). The extent of mesenchymal cell condensation has been related to the level of cell differentiation towards chondrogenesis (Evans and Tuan, 1988; San Antonio and Tuan, 1986).

In the last years, our group has developed a dendrimer-based nanopatterning technique using RGD-functionalized polyamidoamine (PAMAM) G1 dendrimers that allowed us to control the local surface adhesiveness at the nanoscale on large surface areas, thus being compatible with standard cell culture protocols (Casanellas et al., 2018; Lagunas et al., 2014). By using different solution concentrations of the dendrimers, we created different nanopattern configurations by adsorption with a liquid-like order on low-charged surfaces. Although each dendrimer of 4-5 nm in diameter contained eight copies of RGD, due to steric effects, a dendrimer could solely interact with one integrin receptor of around 10 nm in diameter (Lagunas et al., 2014; Xiong et al., 2002). Thereby, dendrimer configuration on the surface matched the adhesive sites for integrin interaction. We produced different nanopatterns on poly(L-lactic acid) and named them according to the percentage of surface area containing inter-dendrimer distances below the 70 nm threshold required for an efficient cell adhesion on stiff surfaces: S_90_ (90%, high adherence), S_45_ (45%, intermediate adherence) and S_18_ (18%, low adherence), respectively (Arnold et al., 2004; Lagunas et al., 2017; Oria et al., 2017). We observed that nanopatterns effectively sustain cell culture for several days and that, under chondrogenic induction, human adipose-derived mesenchymal stem cells (hASCs) formed condensations on all the nanopatterns (Lagunas et al., 2017).

Herein we live-imaged single and collective cell migration dynamics at the early stages of mesenchymal condensation on the nanopatterns. We found that substrate adhesion mediated both single and collective cell movement during condensation under chondrogenic stimuli. On nanopatterned substrates, mesenchymal cells migrate while extending and retracting membrane protrusions to probe both the substrate and other cells and condensates. Whenever two cells come into contact, they can either form stable cell-cell unions, leading to condensation, or eventually detach to resume single cell migration. Successful condensation is therefore dependent on the capacity of cells to migrate, collide, and progressively generate stable multicellular structures. We observed that the cell migration mode on single cells depended on substrate adhesiveness, going from amoeboid-like to more mesenchymal with increasing adhesion. Cells on S_90_ nanopatterns showed traits from both modes of migration, preserving amoeboid features that facilitate migration, but at the same time, enhanced directional movement and larger protrusions that promote condensation. The adhesive characteristics of S_90_ provided an efficient balance between cell-substrate and cell-cell interactions, aiding cell condensation and condensate growth. The anchor points on S_90_ were sufficient to maintain a directional movement and provided the sufficient strength to reduce contact inhibition of locomotion (CIL) events during cell-cell contact, thus favoring condensation. Blockage of cell-cell interactions through *N*-Cad and GJ inhibitors caused a decrease in both the migration rate and directionality in the single cell migration on S_90_ nanopatterns. The directionality of the movement was preserved in the migration dynamics of cell condensates on S_90_, thus favoring their growth through merging. Results from inhibition of cell-cell interactions showed that *N*-Cad-mediated cell cohesiveness is necessary for the directional migration of condensates on S_90_ nanopatterns. Altogether, these results highlight the relevance of the fine tuning of migration characteristics through cell-substrate interactions for an efficient condensation and further chondrogenesis *in vitro*.

## 2. Results

### 2.1. hASCs adapt their migration mode to the adhesive properties of the substrate

hASCs were seeded on the nanopatterned substrates (S_18_, S_45_ and S_90_) in chondrogenic culture medium to induce mesenchymal cell condensation. Pristine non-patterned (S_0_) and fibronectin (FN)-coated substrates (S_FN_) were chosen as representatives of non-adherent (Zhu et al., 2004) and highly adherent surfaces, respectively. We first checked the number of adhered cells in each substrate, which as expected, increased with the RGD ligand density (Fig. 1A). Cell movement was tracked by fluorescence live cell imaging during 40 h. In the absence of symmetry-breaking gradients, mesenchymal cells are expected to migrate randomly (persistent random walks, PRWs) dynamically switching between fast translocation and slow rotation, combining elongated and rounded morphologies, respectively (Wu et al., 2015). We observed that while single cells on S_FN_ substrates adhere and spread within the first 48 min after cell seeding, single cells on the nanopatterns and on S_0_ controls preserved their rounded morphology (Fig. 1B and Supplementary Fig. 1). Differences in cell spreading condition the cell migration mode. Cells on S_FN_ displayed a typical mesenchymal migration, with elongated spindle-like morphologies and a protruding leading-edge, which is rich in actin (Fig. 1C), and cells on S_0_ and the nanopatterns exhibited amoeboid-like migration presenting rounded irregular morphologies that underwent transient expansion and contraction cycles with protruding filopodia (Fig. 1D). Cell spreading also influenced actin network configuration, which was organized in stress-fibers in S_FN_, while single cells on S_0_ and on the nanopatterns displayed a cortical actin organization (Casanellas et al., 2020) that concentrates at the rear part of the cell during migration (Fig. 1E).

**Figure 1.**
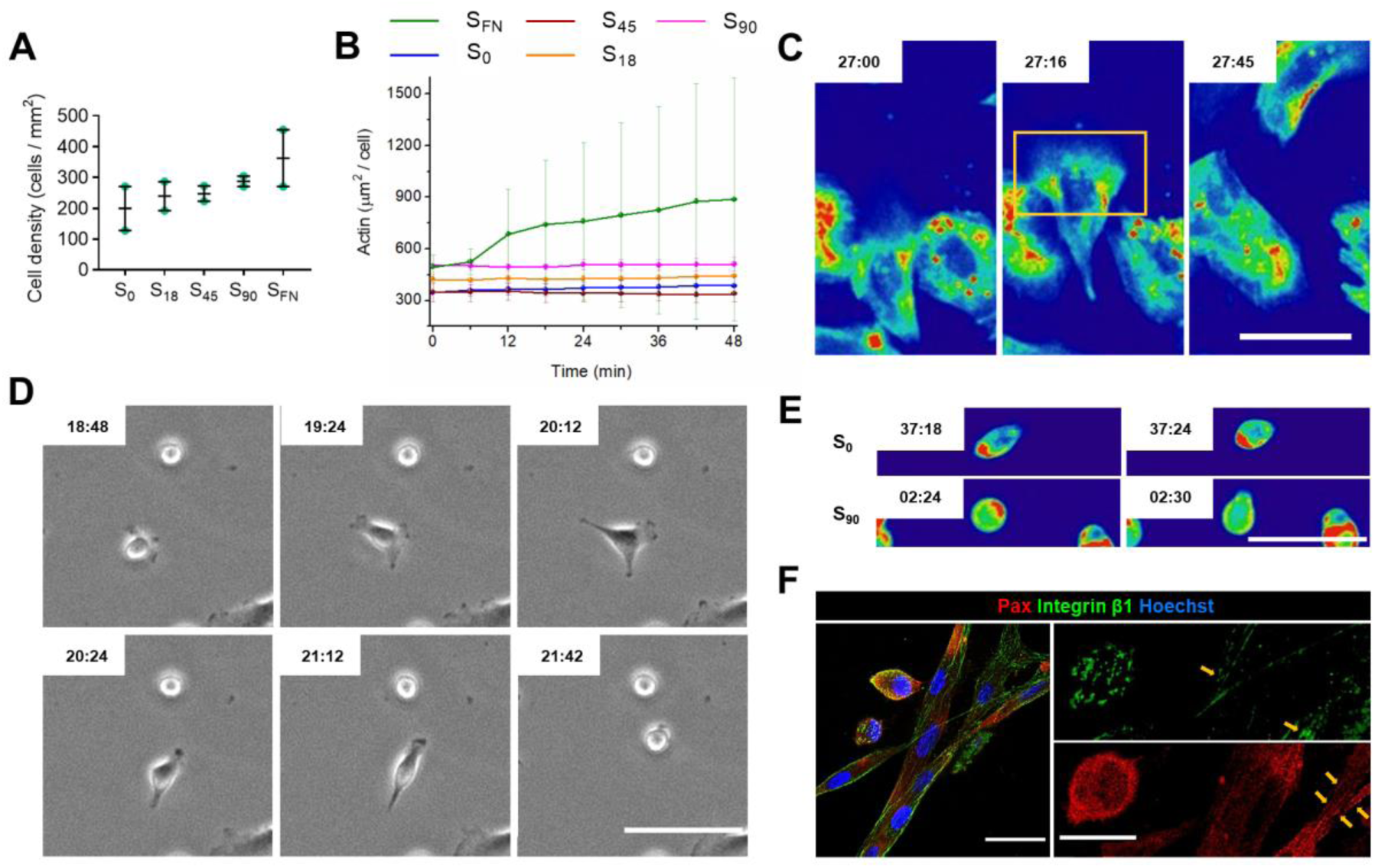
hAMSCs under chondrogenic stimuli adapt their migration mode to the environment. **(A)** Quantification of initial cell density after seeding, 128 ≤ n ≤ 455. **(B)** Quantification of the spreading area from actin staining (SiR-actin) during the first 48 min after cell seeding, n = 9. Results are the mean ± SEM. **(C)** Time-lapse intensity color map for actin staining on S_FN_ showing the actin reach leading front of a mesenchymal migrating cell. Red and blue are the maximum and the minimum, respectively. Scale bar = 75 µm. **(D)** Time-lapse phase-contrast images of an amoeboid migrating cell on S_90_ switching between rounded and extended morphologies. Scale bar = 100 µm. **(E)** Time-lapse intensity color map for actin staining on S_0_ and S_90_ showing cortical actin concentrates at the rear part of the cell during amoeboid migration. Red and blue are the maximum and the minimum, respectively. Scale bar = 80 µm. **(F)** Representative immunofluorescent images of cells on S_90_. Rounded single cells and spread interacting cells can be both observed during cell tracking. Scale bar = 30 µm. Images show that FAs were only present in the spread groups of cells (yellow arrows), while integrin β1 (green) and Pax (red) appeared diffuse within the cytosol in the amoeboid migrating cells. Scale bar = 15 µm.

Amoeboid migration in slow mesenchymal cells has been described for substrate adhesion levels that impede cell binding, and attributed to the absence of focal adhesions (FAs) (Liu et al., 2015). Accordingly, we observed that in amoeboid migrating cells, β1 integrins and the FA adaptor protein paxillin (Pax) were homogeneously distributed and no FAs were formed. Nevertheless, on these samples (both S_0_ and the nanopatterns), Pax-rich FAs could be observed along the perimeter of spreading cells already forming cell aggregates (Fig. 1F), indicating that integrin-based adhesion was not completely inhibited on these substrates. Moreover, the presence of filopodia during transient expansion in amoeboid migrating single cells and in cell-cell interactions, suggested the engagement of integrin receptors (Renkawitz et al., 2009). Quantification of filopodia length on S_0_ and on the nanopatterns demonstrated that cells on S_90_ generate significantly longer protrusions (Supplementary Fig. 2), indicating that cells are sensitive to the RGD integrin ligand density on the substrates. FAs were formed on S_FN_ (Lagunas et al., 2017), thus supporting the traction forces necessary for mesenchymal migration.

### 2.2. Cells migrating on high-adherence substrates show a more directional movement

Quantification of the mean track velocity shows that cells on mid-adherence nanopatterns (S_45_) migrated significantly faster (at around 0.6 µm min^-1^) than on the other substrates (Fig. 2A). This agrees with previous reports showing a biphasic dependence of cell migration velocity, in which a maximal migration rate can be found for an intermediate level of cell-substrate adhesiveness (DiMilla et al., 1993). To quantify directionality, we examined the cell track tortuosity, which was calculated as the relation between the Euclidean distance (the shortest straight distance between starting and end points of the cell’s trajectory) and the total trajectory covered by the cell. A value of tortuosity closer to 1 indicates a more directional movement, whereas lower values reveal a more winding path (values equal to 0 are due to static cells). Tortuosity increased with substrate adhesiveness, and similar values were obtained for S_18_ and S_45_ substrates (Fig. 2B). The more directional migration with substrate adhesiveness could be also qualitatively appreciated when plotting the cell trajectories (Supplementary Fig. 3). Moreover, the turning angles between consecutive frames obtained for cells on S_0_ showed random cell movement, with all angles having a similar probability. With increasing local RGD density, cell motility progressively evolved towards a more directional migration, as shown by the higher proportion of turning angles at low values (Fig. 2C). The calculated mean squared displacement (MSD) of the cell trajectories (Supplementary Methods) also reflected the increase in the directionality of the movement with the adhesion of the substrate. MSD is a measure of the spatial extent of the region explored by the cells. When measured over time, MSD shows a steeper increase with the increasing local ligand density due to the higher directional persistence in cell migration (Fig. 2D). Thereby, our results show that cells on more adherent substrates present a more directional movement that allows them to span over longer distances despite their lower migration rate.

**Figure 2.**
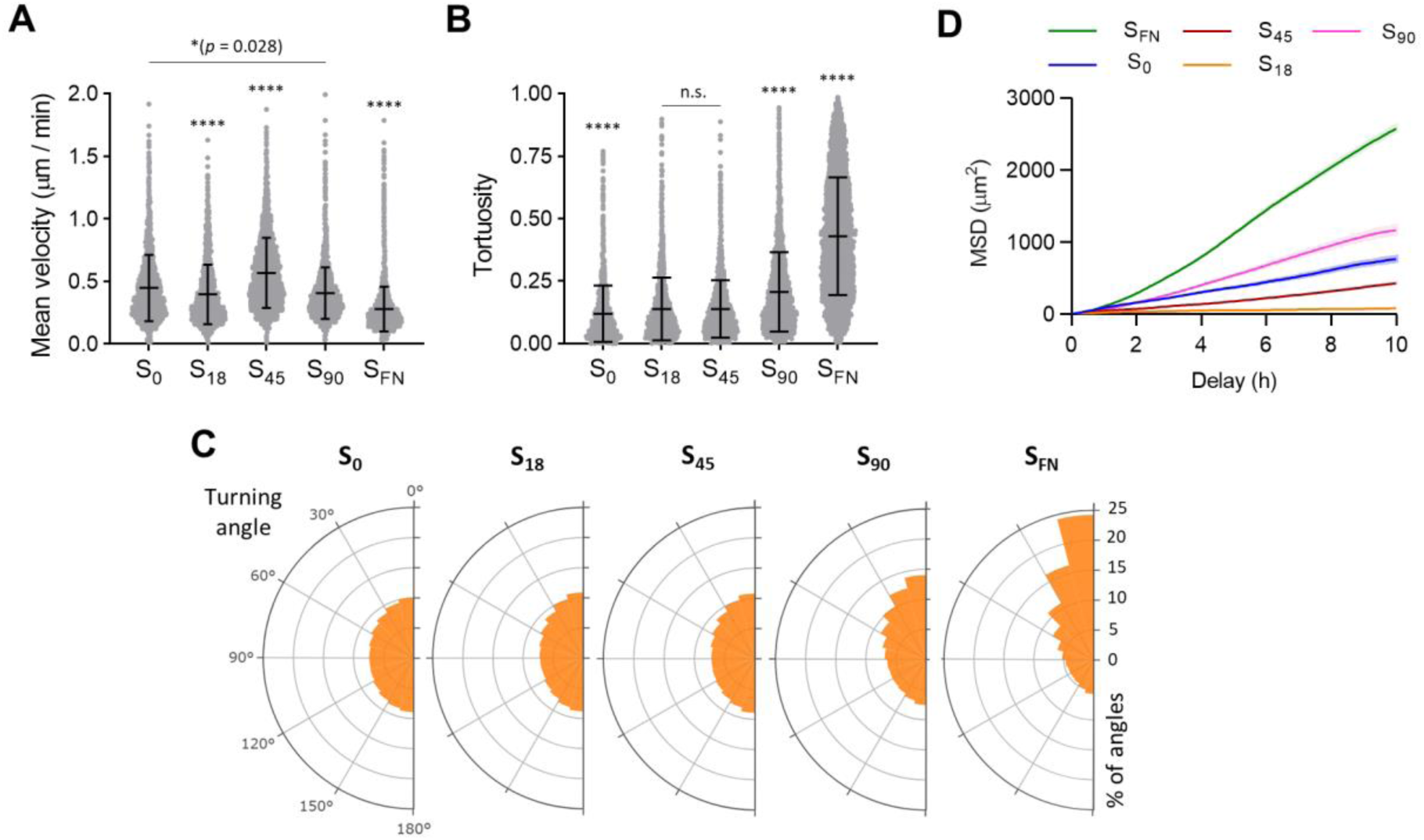
Single cells migrate more directionally with increasing the adhesiveness of the substrate. **(A)** Quantification of mean velocity. Results are the mean ± SD. 2387 ≤ n ≤ 7424. **(B)** Quantification of track tortuosity, and **(C)** Polar histograms of turning angles in single-cell tracks; 309555 ≤ n ≤ 980940. **(D)** MSD for each of the substrates. Results are the mean ± SEM. n = 384. Data are representative of at least two independent experiments with all data from both computed in the calculations. Statistical significance was determined by a multiple-comparison post hoc test. \*\**p* < 0.01, *** *p* < 0.001, \*\*\*\**p* < 0.0001.

### 2.3. Simulated whole-cell migration

To account for cell-substrate interactions, we adapted the model developed by Odde and coworkers of whole-cell simulation of cell migration (Chan and Odde, 2008; Klank et al., 2017). Since each dendrimer of 4-5 nm in diameter is able to bind just one integrin molecule (Lagunas et al., 2014), we modeled the substrate as discrete integrin binding sites defined by the adsorbed dendrimers. The individual integrin-RGD dendrimer bonds and the adaptor proteins were the molecular clutches that link F-actin to the substrate and mechanically resisted myosin-driven F-actin retrograde flow through the stiffness of the clutches and the dendrimer adsorption force (Fig. 3A). In analogy to previous works (Klank et al., 2017), we maintained the number of actin motors constant and we varied the number of clutches in the same proportion in which dendrimers vary on the different nanopatterns, previously determined by atomic force microscopy (AFM) analysis (Lagunas et al., 2017). We set the number of clutches to 20 for S_0_ and set the number of clutches equal to the number of motors for S_FN_. We obtained the simulated cell trajectories for the different substrates (Fig. 3B), from which the MSD and the random motility coefficient (RMC) were calculated (Supplementary methods). As also found by Klank et al., RMC reflected the biphasic dependence of migration speed on the substrate adhesiveness (Klank et al., 2017), peaking in our case at S_45_ in agreement with the experimentally determined mean velocity (Fig. 3C). However, the MSD profiles obtained from the simulation do not reproduce the experimental data (Fig. 3D). Since MSD is a function of both cells mean velocity and persistence (Yeoman and Katira, 2019), discrepancies between the calculated and experimental values could be attributed to the predominant effect of directional persistence, as observed in the experimental data, which lead to deviations from the PRW model.

**Figure 3.**
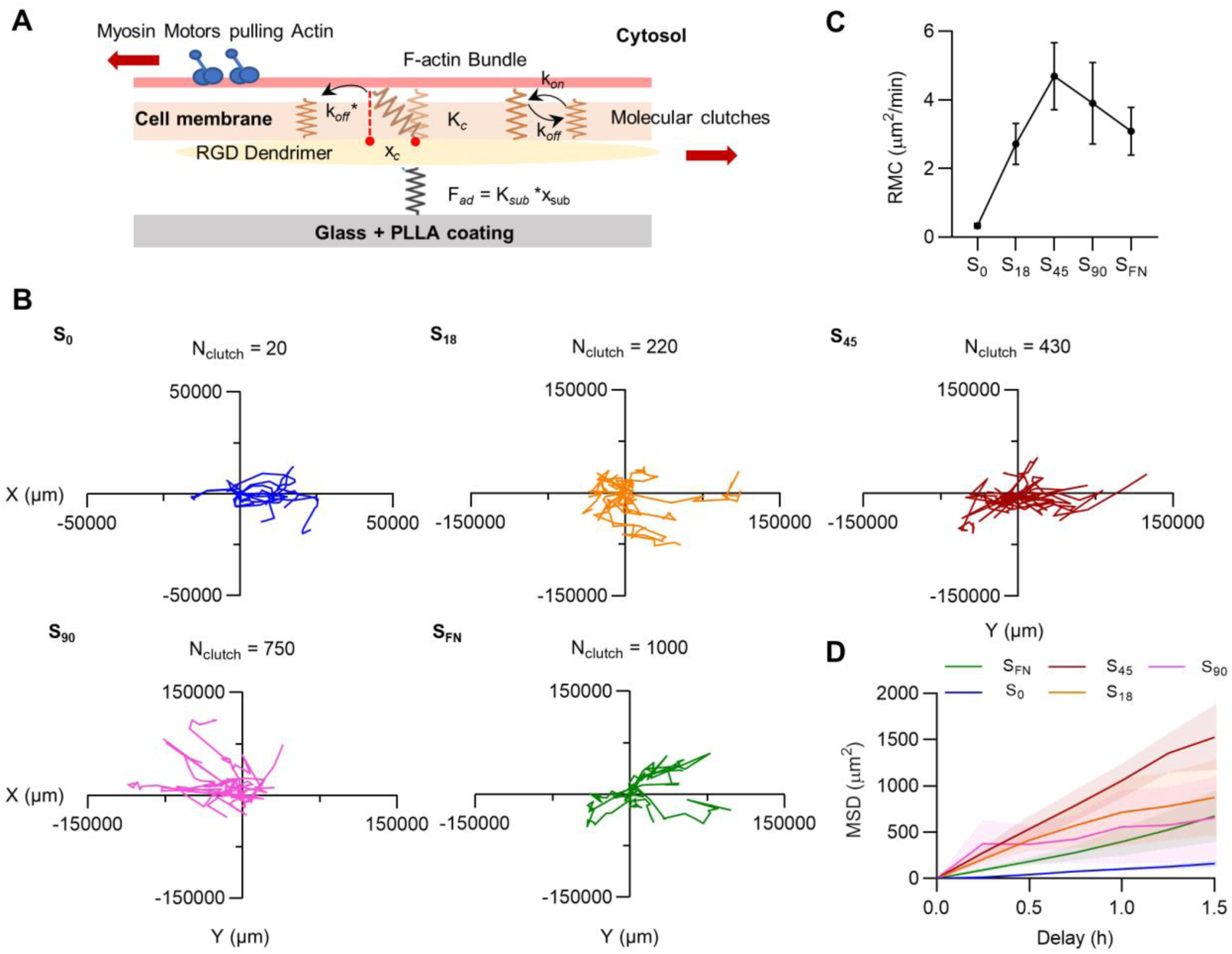
Single cell migration simulation. **(A)** Schematics of a motor-clutch module. Myosin motors pull the F-actin filament bundle retrogradely causing the displacement of the clutches and building tension. The molecular clutches can reversibly engage the F-actin bundle with rates k_*on*_ and k_*off*_ to resist retrograde flow. During loading, the molecular clutches are displaced till they fail at x_c_ with a k_off_* rate, which is force dependent. The mechanical stiffness of the clutches, K_*c*_, and the adsorption force of the dendrimers (F_*ad*_ = K_*sub*_ *x_sub_) will determine the mechanical resistance to loading. **(B)** Plot of representative single-cell trajectories obtained from 5 h simulation and a constant motor number per cell, which was maintained at 1000. The number of clutches (*N*_*c*_) on the nanopatterned substrates was calculated setting 750 clutches for S_90_ and decreasing this number for S_45_ and S_18_ in the same proportion as it does the number of dendrimers inferred from AFM image thresholds (data not shown). *N*_*c*_ was set at 20 for S_0_ and at 1000 for S_FN_. **(C)** Calculated RMC and **(D)** MSD from the simulated cell trajectories. Results are the mean ± SEM.

### 2.4. Cell condensate formation and collective cell migration

Immediately after seeding, cells on S_0_ and on the nanopatterns start aggregating into condensates (Movies 1-4). Conversely, cells on S_FN_ quickly developed a compact monolayer (Movie 5), from which no clusters of cells appeared until day five of chondrogenic induction (Lagunas et al., 2017). During cell condensate formation, cells interacted with each other and generate stable adhesions. The number of cells, velocity, and the directionality of migration in a multicellular environment will affect the number of cell-cell encounters and thus the chances for condensation to ensue. We quantified cell-cell merge events, defined as instances of two tracks reaching a nucleus-nucleus distance lower than 30 µm. The hourly rate of merge events per track was equal or very similar for S_0_, S_18_ and S_90_ (means of 0.22-0.23 events h^-1^), but significantly higher for S_45_ (0.35 events h^-1^) (Fig. 4A). Thereby, suggesting that the number of merge events is facilitated by a higher mean velocity on the substrates, and that there are more chances for cells on S_45_ nanopatterns to initiate condensation.

**Figure 4.**
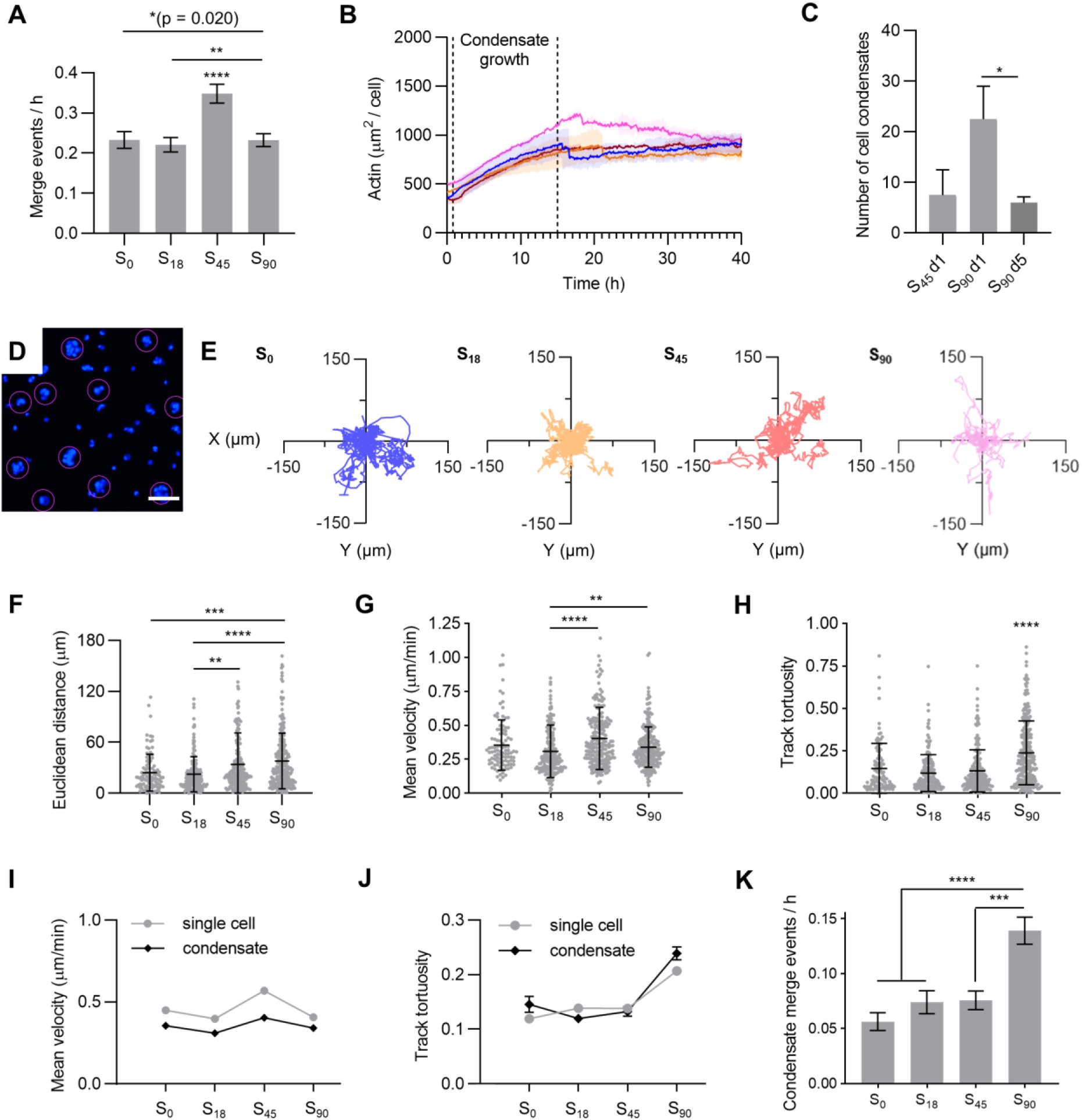
Formation of mesenchymal cell condensates. **(A)** Quantification of the merge events per hour in cell tracks; 941 ≤ n ≥ 2183. **(B)** Quantification of the spreading area from actin staining (SiR-actin) during cell tracking showing the region assigned to cell condensate growth, n = 9. Values from 48 min to 15 h have been fitted into a linear model and results are summarized in Supplementary Table 1. **(C)** Quantification of the number of cell condensates per substrate on S_90_ and S_45_ nanopatterns formed after 1 day of chondrogenic induction; 4 ≤ n ≤ 29. **(D)** Representative image of cell nuclei (blue) showing cell condensates segmented for tracking (contained in purple circles) at 20 h of imaging. Scale bar = 100 µm. **(E)** Migratory trajectories for cell condensates, n = 20. The beginning of each trajectory has been tied to the origin of the Cartesian coordinate system. **(F)** Quantification of the Euclidean distance., **(G)** mean velocity and **(H)** track tortuosity of cell condensates. 101 ≤ n ≤ 247. Results are the mean ± SD. **(I)** Comparison of mean velocity and **(J)** track tortuosity between single cells and cell condensates. 101 ≤ n ≤ 5177. **(K)** Quantification of the merge events per hour in cell condensate tracking. 178 ≤ n ≤ 211. Results are the mean ± SEM. Data are representative of at least two independent experiments with all data from both computed in the calculations. Statistical significance was determined by a multiple-comparison post hoc test. \**p* < 0.05, \*\**p* < 0.01, *** *p* < 0.001, **** *p* <0.0001.

Since during individual cell migration, cells on S_0_ and on the nanopatterns retain their rounded morphology, the increase in the spreading area that occurs during the first 15 h of cell tracking (Fig. 4B, area between dashed lines) can be attributed to cell condensation. Quantification of the spreading area from actin staining shows that the growth rate of the cell condensates, measured as the slope of the curves in Fig. 4B, is faster on S_90_ (see Supplementary Table 1 for fitting parameters and Dataset’s comparison tests). This higher growth rate, which does not correspond with the number of merge events, could be caused either by the formation of more condensates on the sample and/or by condensates growing larger. Previous results showed that S_90_ and S_45_ nanopatterns produced condensates of similar projected areas at day one of culture (Lagunas et al., 2017). Fig. 4C shows that in the same time, more condensates were formed on S_90_ than on S_45_, indicating that merge events on S_90_ nanopatterns, although less frequent, are more efficient in terms of cell condensate formation, whereas on S_45_ a higher proportion of cell encounters did not lead to condensation.

We analyzed the migration dynamics of multicellular cell condensates on the substrates. In analogy to single cell migration analysis, whole condensates were segmented and tracked (Fig. 4D), starting at 20 hours of imaging. Cell condensate trajectories are shown in Fig. 4E. We found that substrate adhesiveness had similar effects on collective cell migration as on single cells. The quantification of the Euclidean distance shows an increase with substrate adhesiveness (Fig. 4F), which is attributed to a more directional cell movement, supported by the mean velocity and tortuosity results (Figs 4G and 4H, respectively). Cell condensates on S_45_ and S_90_ nanopatterns had a similar migration rate (of around 0.4 µm min^-1^), whereas the directionality of the movement was significantly higher on S_90_, as also appreciated in the quantification of the turning angles (Supplementary Fig. 4). The rate of migration was significantly slower for cell condensates on all the substrates compared to single-cell migration, although the shape of the curve of the mean velocity against substrate adhesiveness was preserved (Fig. 4I). Track tortuosity values (Fig. 4J) show that the directionality of the movement was very similar in single-cell migration and in condensate migration.

During live imaging, we observed several instances of two or more cell condensates coming into contact and then fusing through a coalescence-like mechanism, leading to the formation of bigger structures (Supplementary Movie 1a and b). We previously observed that the size of the cell condensates increases during the first five days of chondrogenic induction, which can be produced by the incorporation of cells to an existing cell condensate and/or by merging of cell condensates. On S_90_nanopatterns, there is a significant decrease in the number of cell condensates from day 1 to 5 of culture (Fig. 4C). Considering that on S_90_ nanopatterns the condensates are stable for up to 14 days in culture (Casanellas et al., 2020), we attributed the decrease in their number to the occurrence of merging events, which were about two times more frequent on S_90_ nanopatterns than on the rest of the substrates (Fig. 4K).

### 2.5. Blocking cell-substrate and cell-cell interactions

Tracking results suggest that substrate adhesion on S_90_ nanopatterns induces an intermediate mode of migration, where cells exhibit most common amoeboid traits but present higher directional persistence, which could be advantageous for cell condensate formation and growth. We analyzed migration and condensation under three types of pharmacological interventions on these substrates impairing certain cell-substrate and cell-cell interactions: in-solution RGD dendrimers, ADH1 peptides and 18β-glycyrrhetinic acid (18βGA). Addition of RGD peptides in culture medium impairs integrin clustering (Cluzel et al., 2005; Wang et al., 2010); we thus expected that RGD dendrimers in solution would have a similar effect, hampering integrin interaction with substrate-adsorbed ligands by competition. ADH1 is a cyclic pentapeptide that contains the HAV sequence that binds to adherens junction protein *N*-Cad (Erez et al., 2004; Williams et al., 2000), thus blocking it when added in cell culture (Madl et al., 2017; Madl et al., 2019; Shintani et al., 2008). Phosphorylation of gap junctional connexin proteins regulates several aspects of gap junction intercellular communication (GJIC), including GJ formation, gating and turnover (Pogoda et al., 2016; Solan and Lampe, 2014; Solan and Lampe, 2018). 18βGA is a saponin that induces disassembly of GJ plaques through connexin dephosphorylation (Böhmer et al., 2001; Guan et al., 1996).

We imaged single cell and multicellular cluster dynamics on S_90_ substrates under each of the pharmacological treatments (Movies 6-8). All treatments caused a significant reduction in the directional persistence of the single cell movement, although integrin blockage had a stronger effect: a 2-fold decrease in tortuosity was measured with the addition of RGD dendrimers in solution (Fig. 5A). This indicates that cell adhesion to the substrate is more relevant than cell-cell interactions to determine the directionality of single cell migration. Moreover, the value of tortuosity after treatment was even lower that the one measured on S_0_ control substrates (Fig. 5B). Quantification of the mean cell velocity showed that all pharmacological treatments caused a significant, although small, reduction in the migration speed. Integrin blocking and the uncoupling of GJs with 18βGA had a similar effect (cells were around 25% slower) and blocking *N*-Cad with ADH1 reduced the mean velocity of the cells by 13% (Fig. 5C). The merge events per hour were significantly reduced with all the treatments, especially with integrin blockage. Both ADH1 and 18βGA treatments reduced the hourly rate of merge events between tracks by 40%, while integrin blocking induced a maximum reduction of 55% (Fig. 5D).

**Figure 5.**
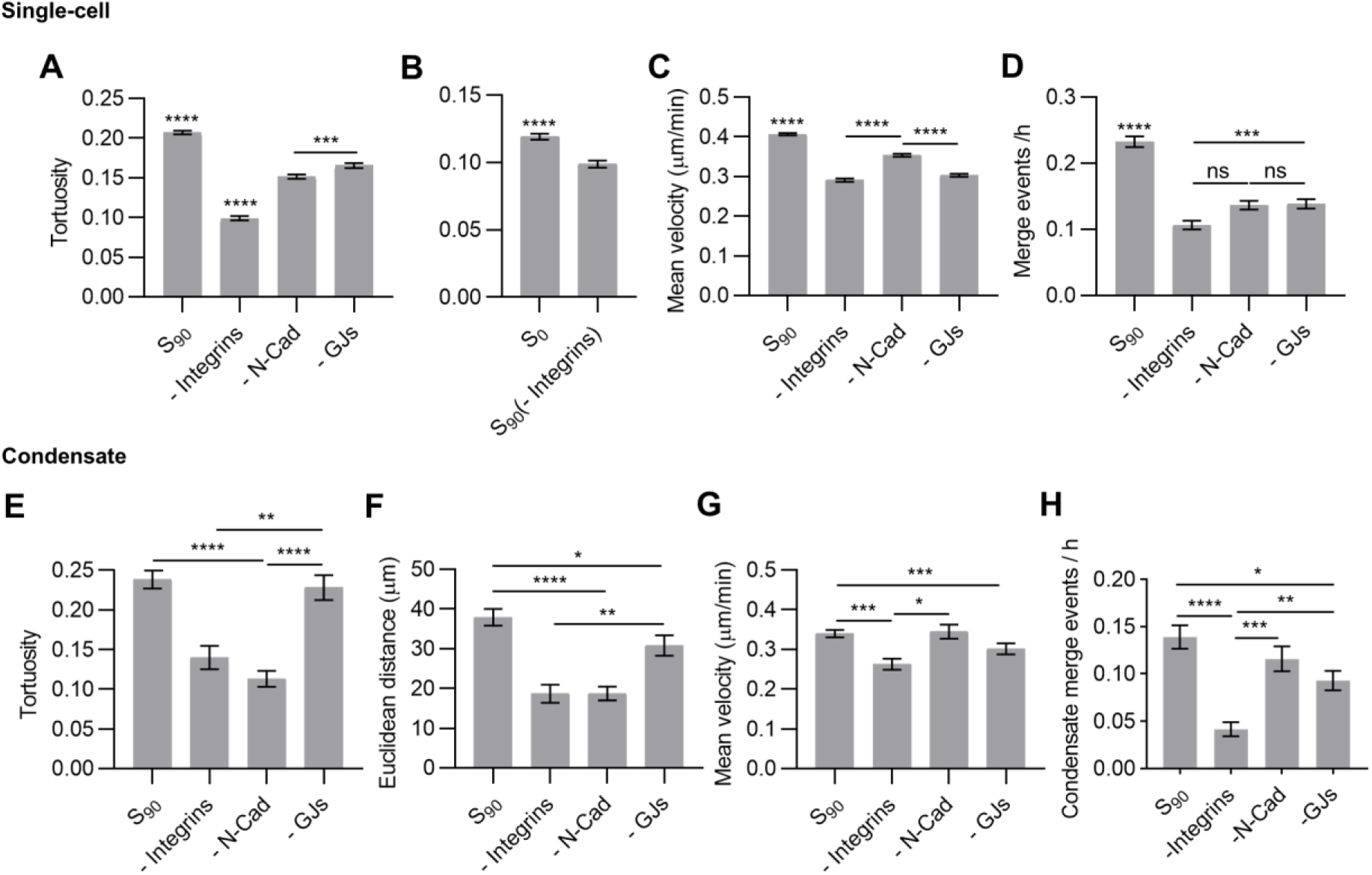
Effects of pharmacological inhibitors on single cell and in collective cell migration on S_90_ nanopatterns. **(A) and (B)** Quantification of tortuosity, **(C)** mean velocity, and **(D)** merge events per hour in single cell tracking. 1574 ≤ n ≤ 5177. **(E)** Quantification of tortuosity, **(F)** Euclidean distance, **(G)** mean velocity and **(H)** merge events per hour in collective cell migration of cell condensates. 85 ≤ n ≤ 247. Results are the mean ± SEM from S_90_ without (S_90_) and with the pharmacological interventions: RGD dendrimers in solution (-Integrins), ADH1 (-*N*-Cad), and 18βGA (-GJs), respectively. Data are representative of at least two independent experiments with all data from both computed in the calculations. Statistical significance was determined by a multiple-comparison post hoc test. \**p*<0.05, \*\**p*<0.01, \*\*\**p*<0.001, \*\*\*\**p*<0.0001.

In the analysis of cell condensates migration, we observed that directionality is affected by integrin blockage causing a 41% decrease in tortuosity, and by *N*-Cad inhibition with a 53% decrease (Fig. 5E). However, GJ disassembly had a lesser (19% decrease) to no significant effect on directional persistence (Figs 5F and 5E). Similar to single cell tracking results, quantification of the mean velocity in cell condensates (Fig. 5G) shows that GJ disassembly and integrin blockage cause a significant, but small, reduction of the migration rate (11% and 23% decrease, respectively), and no significant effects were observed upon *N*-Cad inhibition. The merge events between cell condensates were severely affected by integrin blockage, which caused a 3-fold decrease in their hourly rate, whereas the disassembly of GJs caused a 33% decrease and *N*-Cad inhibition had no significant effects.

## 3. Discussion

Cells cultured on the substrates show a range of traits that go from more amoeboid to more mesenchymal with increasing substrate adhesiveness. Cells on S_FN_ controls spread quickly after seeding, displaying a typical mesenchymal migration with spindle-like elongated morphologies and exerting traction forces against the substrate through FAs associated with actin-rich protrusions. Cells on S_FN_ are the slowest of all analyzed conditions, in agreement with previous reports indicating that the velocity of amoeboid migration is usually higher than that of mesenchymal migration (Panková et al., 2010; Ruprecht et al., 2015).

Conversely, cells on S_0_ and on the nanopatterns exhibited amoeboid-like migration with rounded irregular cell morphologies and transient expansion and contraction cycles with protruding filopodia. They show cortical actin organization that concentrates at the rear part of the cell during migration, which has been reported to be a consequence of the localized retrograde flow that generates by active actin polymerization in the filopodia and pushes the cortex backwards (Liu et al., 2015; Renkawitz et al., 2009). Amoeboid migration is generally observed in cells migrating on low-adherence substrates under spatial constraints, such as in microfluidic devices or in hydrogels (Liu et al., 2015). It has been recently reported in cancer and epithelial cells migrating freely on soft, viscoelastic substrates (Adebowale et al., 2021), as well as neural progenitors cultured at low density (Kawaguchi et al., 2017).

Cells can switch between migration modes depending on the context (Yamada and Sixt, 2019). Migration plasticity between the mesenchymal and amoeboid migration has been extensively studied in cancer dissemination, where it is termed mesenchymal-to-amoeboid transition (MAT). MAT endows mesenchymal cells with amoeboid features that facilitate migration through complex 3D environments, aiding cancer metastasis (Graziani et al., 2022). The switching between migration modes has been related to three key parameters: protrusion, contractility, and adhesion (Lämmermann and Sixt, 2009), and the transition to fast amoeboid-like migration has been described under conditions of low adhesion and strong confinement (Liu et al., 2015). Here, we observed that hASCs under chondrogenic induction adapt their migration mode to the substrate adhesiveness, influencing both single cell and collective cell migration during cell condensation.

Significantly longer protrusions were generated by cells on S_90_ nanopatterns, indicating that they are sensitive to the RGD integrin ligand density. Although no FAs were present, amoeboid cells migrating on 2D substrates require the surface receptors to adhere to the substrate to prevent Brownian motion from keeping them apart of surface contact, and to sustain the actin retrograde flow during migration. We observed that β1 integrins were homogeneously distributed on the plasma membrane of the cells, as described previously for amoeboid T lymphocytes (Friedl et al., 1998). These diffusively distributed anchor points have been found to be sufficient to generate the traction forces necessary for the migration of low-adhesive cells (Graziani et al., 2022; Lämmermann and Sixt, 2009).

Quantification of the mean velocity from cell tracking showed that cells migrated significantly faster on mid-adherence substrates (S_45_) in agreement with previous studies demonstrating a biphasic dependence of cell migration velocity with cell-substrate adhesiveness, where maximum speed is reached at intermediate levels of adherence (DiMilla et al., 1993). Cell migration based on anchoring receptors requires de dynamic building and breakage of attachments with the substrate. On low adherent substrates (S_0_ and S_18_), the scarceness of anchor points compromises traction force during migration, whereas with the increasing substrate adherence, transient contacts become more stable, enhancing traction force but also making rearward detachment more difficult. A balance between these two situations is given by S_45_ nanopatterns, thus inducing a peak in the migration rate.

We examined three different indicators of directional persistence: tortuosity, turning angle and MSD. MSD contains information about both speed and directional persistence and is a good measure of the surface area explored by the cells over time. We found that higher substrate adhesiveness makes the cell movement more directional. Directional persistence in cell motility relies on preserving the established front-rear polarity upon cell symmetry breaking (Hennig et al., 2020). The increasing availability of adhesion sites (higher adhesiveness) could minimize turning, and thus favor persistence. However, at the same time, it might introduce conflicting directional signals. Sung et al. demonstrated that the autocrine secretion of exosomes from late endosomes is required for directionally persistent cell movement of migrating cancer cells *in vivo* and *in vitro*. Exosome secretion precedes and promotes adhesion assembly at the leading edge, stabilizing nascent protrusions, which become larger and long-lived. *In vitro* experiments show that FN from serum or from surface coatings is targeted to exosomes by binding to cognate cellular integrin receptors, and then secreted in adhesive form to promote adhesion formation (Sung et al., 2015). Although most probably also aided by other signaling cargos (Luga et al., 2012), adhesive ligand availability (RGD or FN) in most adherent substrates (S_90_ and S_FN_, respectively) might favor adhesion recycling, exosome formation and trafficking, thereby promoting polarization maintenance and directionality. However, the abrupt increase of directionality on S_FN_ substrates (2-fold increase in tortuosity compared to S_90_) could not be explained in terms of ligand density alone, as S_90_ and S_FN_ substrates present a relatively small difference in the percentage of area covered with RGD groups at less than 70 nm: from 90% for S_90_ to presumably 100% for S_FN_, given a molecule size of 15.5 × 8.8 nm (Koteliansky et al., 1980). However, the spatial positioning of RGD groups on native FN would differ from that of nanopatterned RGD dendrimers, generating different sets of cell-substrate interactions and differently affecting cell adhesion and migration (Cavalcanti-Adam et al., 2007). Moreover, integrin binding to fibronectin is also modulated by the synergy sites at the protein, which are absent in isolated RGD dendrimers (Brown et al., 2015). Finally, a coating of whole fibronectin protein does not engage all the same integrin types that isolated RGD motifs do (Humphries et al., 2006; Moreno-Layseca et al., 2019), which can also contribute to a difference in cell behavior if the activated integrins have different parallel biological functions (Elosegui-Artola et al., 2014).

Our results point to the importance of directionality, rather than speed, to determine how far a cell will migrate. While moving at greater speed allows cells on S_45_ to cover a longer sum trajectory, their lack of directionality means that the Euclidean distance remains lower than those of S_90_ and S_FN_ (Supplementary Fig. 5). On the other hand, moving in straighter paths allows cells on S_FN_ to migrate farther and explore larger surface areas as demonstrated by MSD measurements, even with a lower velocity than all other conditions. Tellingly, S_0_, S_18_ and S_90_ have very similar mean velocities (lower than S_45_); nevertheless, a more directed movement also allows S_90_ to migrate slightly farther than S_0_, S_18_ and S_45_.

To account for cell-substrate interactions, we performed a whole-cell simulation of cell migration by adapting the model developed by Odde and coworkers (Chan and Odde, 2008). We found that calculation of RMC shows a biphasic dependence with adhesion as previously reported (Klank et al., 2017), which peaks at S_45_ effectively reproducing the experimental data. However, we found discrepancies between empirical and simulated MSD curves that we attributed to variations on the directional persistence among substrates. Indeed, deviations from the PRW model have been reported, especially at large time intervals (Dickinson and Tranquillo, 1993; Dieterich et al., 2008). Alternatively, other mechanisms have been proposed such as the zig-zagging observed in the motile amoeba *D. discoideum*, which aid to maintain directional persistence during cell movement (Li et al., 2008).

The chondrogenic cell culture medium induces cell aggregation on the substrates into cell condensates (Lagunas et al., 2017). During condensation, cells gather and establish intimate cell-cell contacts, for which migrating cells need to collide and become engaged. To evaluate cell condensate formation, we first quantified the rate of merge events as the instances of two tracks reaching a spacing of less than 30 µm between the nuclei of migrating cells, at which we assume cell-cell contact is established. We found that the probability of cell-cell encounters is higher for the fastest migrating cells (those on S_45_ nanopatterns). It should be noted that condensation involves not only merging, but also splitting events in which cells that established transient adhesions come apart. In live imaging, single cells sometimes probe a condensate and join it for only a few minutes before exiting; other times, a cell is pulled from a condensate by those in another one nearby. Hence, while the rate of merge events does not directly correlate with the number of cells present in the final constructs, it reveals S_45_ as a more dynamic system, in which cells come into contact more often than those on other substrates. However, although more frequent, cell-cell collisions on S_45_ lead to fewer condensations than on S_90_ nanopatterns.

Cell-cell engagement during migration is restricted by events of contact inhibition of locomotion (CIL). CIL is a process that was first proposed by Abercrombie and Heaysman through which a cell ceases motility or changes its trajectory upon collision with another cell. Cells exhibiting CIL do not crawl over each other (Abercrombie and Heaysman, 1954). As suggested previously, CIL-mediated repulsion shall be reduced or switched off during mesenchymal cell condensation in chondrogenesis to facilitate cell-cell interactions (Stramer and Mayor, 2017). In neural crest cells, cell-matrix adhesions are disassembled near the cell-cell contact immediately after collision, which generates the tension across the cell contact needed to induce separation during CIL (Roycroft et al., 2018). As fewer cell-substrate adhesions are built on S_45_ than on S_90_ (due to fewer anchor points being available), we assume that their disassembly at the cell-cell contact will be energetically more favorable, thus promoting CIL and preventing condensation. Conversely, inhibition of CIL-mediated repulsion resulted in an increase in the length, area and stability of cell-matrix adhesions together with a change in actin behavior, which organizes in stress fibers near the contact (Roycroft et al., 2018). This could explain the observation of Pax-rich FAs on those cells already forming cell condensates on S_0_ substrates and on the nanopatterns. The disassembly of cell-matrix adhesions at the cell-cell contact is reported to be mediated by the focal adhesion kinase (FAK)/Proto-oncogene tyrosine-protein kinase (Src) signaling that works downstream of *N*-Cad. Upon contact, cells establish a functional *N*-Cad-based adhesion that activates Src. Src, through FAK, promotes the disassembly of cell-matrix adhesions at the cell-cell contact, reducing traction and increasing tension, which is required for cell separation during CIL (Roycroft et al., 2018).

Mesenchymal collective migration is characterized by the cooperative interaction of groups of cells that can be connected either through transient or stable cell-cell contacts, depending on cell density (Theveneau and Mayor, 2013). We analyzed the migration dynamics of condensates on the substrates, and we found that substrate adhesiveness has comparable effects on collective cell migration and on single cells. Similar migration behaviors in individual and collective groups of migrating cells have been previously reported. Mesodermal cells isolated from the migrating cohort were able to migrate independently and in a similar way to the whole group (Arboleda-Estudillo et al., 2010; Ulrich et al., 2005), and in a more recent study, mouse cancer cells forming multicellular spheroids on 2D substrates followed the same migration modes as single cells (Beaune et al., 2018). Our results show that the rate of migration was similarly reduced for cell condensates on the different substrates if compared to single-cell migration, thus indicating that cell condensates are slower than single cells independently of the substrate adherence. We also observed that the directionality of the movement was preserved from single cells to cell condensates on the same substrates. Like in single-cell migration, cell condensates must distinguish a front and a rear to migrate directionally and maintain a certain degree of cohesiveness among the cell cohort to coordinate the movement. As in other models of mesenchymal collective migration (Scarpa and Mayor, 2016), we observed that the whole cell condensate is migratory with cells rapidly exchanging positions and forming large lamellipodial protrusions in the direction of migration at the edge of the condensate (Supplementary Movie 2). These protrusions, in analogy to single migrating cells, exert traction forces against the ECM to allow collective cell movement. As observed in *Xenopus* mesendoderm cells and neural crest cells, integrin-FN interactions are required for migration and deployment into the tissue (Alfandari et al., 2003; Boucaut and Darribere, 1983). The inhibition of α5β1 integrin impairs mesendoderm migration and protrusion formation (Davidson et al., 2002). This agrees with our observation of integrin blockage having a major impact on the migration of both single cells and mesenchymal cell condensates on S_90_ nanopatterns, especially affecting the directionality of the movement, which reaches values below those on S_0_ controls. Indeed, during mesenchymal development cell-substrate interactions become progressively more relevant, in an environment in which the matrix becomes denser over time (Kalson et al., 2015; Singh and Schwarzbauer, 2012).

Alongside cell-substrate interactions, cell-cell adhesions and cell communication have been shown to modulate the migration speed and directionality of collectively migrating mesenchymal cells. *N*-Cad is the main adherens junction protein, mediating cell-cell interactions during mesenchymal cell condensation, and previous studies have shown that the expression of mutant forms of *N*-Cad, lacking either the extracellular homotypic interaction domains or the intracellular β-catenin binding site, resulted in impaired condensation and chondrogenesis (DeLise and Tuan, 2002; Oberlender and Tuan, 1994). Knockdown *N*-Cad in bone marrow-derived mesenchymal stem cells compromised migration in response to transforming growth factor *β* from breast tumor, and *N*-Cad inhibition decreases colony spreading and average spreading velocities in a bladder cancer cell line (Zisis et al., 2022). Accordingly, we found that blocking *N*-Cad with ADH1 reduced the mean velocity of the single cells on S_90_ nanopatterns. These results are consistent with single cell migration within a multicellular setting being mediated by the transient contacts that occur between mesenchymal cells (Theveneau and Mayor, 2012). Strikingly, while no significant effects were found by inhibiting *N*-Cad in the velocity of cell condensates, the tortuosity was decreased at similar levels than for integrin blockage. This magnification of the effects of *N*-Cad on the directionality of the movement in cell condensates can be attributed to its role in maintaining the group cohesiveness during migration.

Mesenchymal cell condensation proceeds with the formation of a widespread GJ communication network that allows cell synchronization and mediates the coordination within multicellular groups towards the formation of tissue (Coelho and Kosher, 1991; Mayan et al., 2015). Connexin 43 (Cx43)-based GJs are essential in chondrocyte differentiation and its deficiency led to skeletal defects (Gago-Fuentes et al., 2016). We found that uncoupling of GJs with 18βGA caused a reduction in the tortuosity and mean velocity of single cells migrating on S_90_ nanopatterns. This is consistent with previous single cell tracking data of mesenchymal cells from elongating feather bud of chicken dorsal skin explants. Authors found that GJ-electric coupling mediates the conversion of sporadic Ca^2+^ transients into organized oscillations that facilitate collective mesenchymal cell migration in feather elongation in the posterior-distal direction. Treatment with the inhibitor carbenoxolone (an 18αGA derivative) was found to alter cell movement, especially the directionality (Li et al., 2018). We found that treatment of 18βGA on migrating mesenchymal cell condensates on S_90_ nanopatterns caused a slight decrease in the migration rate and surprisingly, no significant effects on tortuosity. Li et al. suggested that the mechanism they described involving GJs in the collective mesenchymal migration in feather elongation is active in transient cell collectives to aid them to coordinate during the formation of organ architectures, and it might not be required in more stable adhesion molecule mediated cell condensates.

Although merge events in single cell migration seemed to be dictated by the mean velocity, with cells on S_45_ substrates showing the highest migration rate and producing the highest number of hourly merge events, the dramatic impact of the pharmacological interventions on the merge events per hour in cells on S_90_ nanopatterns, which showed a major decrease in tortuosity compared to the migration rate after treatment, suggested that the directionality of the movement has a non-negligible effect on cell-cell collisions. Importantly, the merging events per hour in cell condensates were found significantly higher on S_90_ substrates, where the groups of cells move more directionally.

We have shown previously that mesenchymal cell condensates produced on S_90_ nanopatterns were mechanically more stable than those produced on the other substrates, preserving the 3D configuration for up to 14 days in culture, and were the only ones able to enter the early stages of chondrogenic pathway, after six days of chondrogenic induction (Casanellas et al., 2020; Lagunas et al., 2017). To avoid hypertrophy as the most probable final outcome of mesenchymal stem cells stimulated towards chondrogenesis, current regenerative medicine approaches focus on controlling the most primary stages of the chondrogenic commitment to favor the formation of hyaline cartilage (Chen et al., 2015). In that sense, it seems a natural approach to tackle substrate adhesions during cell gathering and condensate formation as one of the first steps in cartilage development. Here we show that substrate adhesions have a dramatic effect in both single cell and collective migration of condensates. Single cells migrating on S_90_ nanopatterns present characteristics that could be considered as intermediate between amoeboid and mesenchymal migration, which favor both cell gathering and condensation. Cells show increased directionality and reduced CIL, allowing them to condensate, but at the same time providing a sufficiently dynamic framework in which there is a balance between cell-substrate and cell-cell adhesions for condensates to grow. Therefore, we envision that the results obtained could have an immediate application in cartilage regeneration strategies, but could also be applied to the study of other biological processes involving transitions between the mesenchymal and amoeboid migration modes, such in cancer progression.

## 4. Methods

### 4.1. Preparation of PLLA-coated substrates

Nanopatterned substrates were prepared as previously described(Casanellas et al., 2018; Lagunas et al., 2017). Corning® glass microscopy slides (Sigma-Aldrich, Madrid, Spain) were cut with a diamond-tip cutter to square pieces of 1.25 × 1.25 cm. Slides were washed thoroughly with deionized water (18 MΩ·cm Milli-Q, Millipore, Madrid, Spain) followed by 96% ethanol and air-dried. A 2% m v^-1^ solution of 95/5 L-lactide/DL-lactide copolymer (PLLA, Corbion, Barcelona, Spain) was prepared by adding 200 mf of solid polymer to 10 ml of 1,4-dioxane (Sigma-Aldrich, 296309, Madrid, Spain) in a pressure tube with a magnetic stirring bar. The tube was placed in a silicon oil (Thermo Fisher Scientific, 174665000, Barcelona, Spain) bath on a hot plate at 60°C with gentle stirring (300 rpm) overnight, and the solution was transferred to a 15-ml vial for storage at room temperature. Glass substrates were spin-coated with the PLLA solution in a class 10,000 clean room. Slides were placed on a hot plate at 60°C for at least 10 min to dry. Each slide was fixed on the spin-coater with vacuum and 200-250 µl of PLLA solution were added with a Pasteur pipette, covering the whole surface. Slides were coated with a two-step program: 5 s at 500 rpm with an acceleration of 300 rpm s^-1^ (to eliminate excess solution and spread the remaining solution homogeneously on the surface) followed by 30 s at 3000 rpm with an acceleration of 1500 rpm s^-1^.

### 4.2. Nanopatterning of RGD-Cys-D1 dendrimers on substrates

All steps were performed in a sterile tissue culture hood, and only sterile materials, solutions and techniques were used. Spin-coated PLLA substrates were treated for 13 min under UV light. Deionized water was used to prepare RGD-functionalized dendrimer solutions. A stock solution was prepared by dissolving the solid dendrimer in water, an intermediate solution and three working solutions were prepared from it. Dendrimer solutions were filtered through a 0.22 μm Millex RB sterile syringe filter (Merck Millipore, Madrid, Spain) attached to a syringe and applied directly on the substrates. Substrates were left at room temperature overnight (16 h). Solutions were removed with a pipette in a cell culture hood. Substrates were washed with sterile deionized water and left to dry on air. Fibronectin-coated substrates (S_FN_) were produced by incubating PLLA substrates in fibronectin from bovine plasma solution (Sigma-Aldrich, F1141, Madrid, Spain) 100 μg ml^-1^ in PBS for 1 h at room temperature, and washing with PBS, just before cell seeding.

### 4.3. Cell culture

Human adipose-derived MSCs (ATCC, PCS-500-01, VA, USA) were cultured in T75 flasks at 37°C and 5% CO_2_ in growth medium, consistent of MSC Basal Medium (ATCC, PCS-500-030, VA, USA) supplemented with MSC Growth Kit Low Serum (ATCC, PCS-500-040, VA, USA) and 0.1% v/v penicillin-streptomycin (P/S, Invitrogen, 15140, Barcelona, Spain). Medium was replaced every 2-3 days. Passaging was carried out when cells reached 70-80% confluence.

### 4.4. Live imaging

Prior to imaging, hMSCs at passage 3-4 were incubated in growth medium with SiR-actin (Tebu-bio, SC001, Barcelona, Spain) at 200 nM for 4 h, or 750 nM for 1.5 h. Cells were then incubated in medium with Hoechst (Invitrogen, H3570, Barcelona, Spain) 1:1000 for 5 min. Cells were trypsinized and seeded on the substrate at a density of 30,000 cells cm^-2^ in a glass-bottom microscopy dish (VWR, 734-2906H, Barcelona, Spain) in chondrogenesis-inducing medium (ATCC, PCS-500-051, VA, USA) with SiR-actin 100 nM. The sample was immediately transferred to the microscope setting, pre-conditioned to 37°C and 5% CO_2_. Live imaging was performed in a Nikon Eclipse Ti2-E inverted microscope with a Prime 95B sCMOS camera (Photometrics, Birmingham, UK) and an Okolab Cage Incubator (Okolab, Pozzuoli, Italy), with a 10X objective. Samples were imaged for 40 hours. Images were taken every 6 minutes for the phase contrast, blue and far red (Cy5) channels. At least two samples of each condition were imaged and analyzed.

### 4.5. Single and collective cell segmentation and tracking

Live imaging stacks of Hoechst staining were analyzed for cell movement. The first 17 frames (96 minutes) were removed to omit drifting prior to cell adhesion. Cell nuclei were tracked with the TrackMate plugin in ImageJ (ImageJ version 2.0.0, MD, USA). Nuclei were segmented with the LoG Detector setting an estimated blob diameter of 15 µm and a threshold of 4 µm. No further filters were applied. The LAP Tracker was used with a maximum frame-to-frame linking distance of 50 µm and allowing gap closing to maximum 50 µm over 1 frame. For analysis, frame depth was not limited.

Analysis of collective cell migration in multicellular condensates was performed on the second half (20 to 40 hours) of live imaging Hoechst stacks as described above for single cells, except for the following parameters: Condensates were segmented with an estimated blob diameter of 70 µm and a threshold of 40 µm, and gap closing was allowed over 3 frames. All subsequent steps in track analysis were performed in the same way for single cells and condensates.

### 4.6. Track analysis

Tracks with duration under 3 hours were excluded to avoid skewed results due to artificially short tracks. However, analysis of all available tracks rendered highly similar results.

Track net displacement and duration were extracted from the Track Statistics output file of TrackMate. Sum trajectory was calculated with R Studio (R Studio version 4.0.2, MA, USA) from the Links Statistics output file by adding the displacements of all links in each track. Mean track velocity was calculated by dividing track trajectory (in µm) by track duration (in minutes).

To analyze track directionality, tortuosity was first obtained as the division between the net displacement and the sum trajectory of each track. Turning angles were calculated on Excel from the Spots Statistics output of TrackMate. Vector components were found by subtracting particle coordinates at the start of each 6-minute link from those at the start of the next one (two consecutive spots) within each track. The angles between pairs of consecutive vectors were then found as:

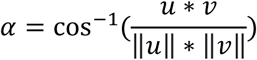

**Equation 1**. Angle α between vectors u and v.

Angles with values equal to 0 or 180 were excluded to account for static cells and single-link tracks. Angles were categorized into 15-degree sections according to their value, section percentages were calculated and plotted in polar histograms with the Plotly package in R Studio (R Studio version 4.0.2, MA, USA).

The number of merge events per track was obtained by running a second TrackMate analysis with the settings as above but allowing track merging when nuclei collided at a distance under 30 µm, for single cell migration; or under 120 µm, for multicellular condensates. The number of merge events per tracks was divided by the duration to obtain an hourly rate of merge events.

### 4.7. Actin staining and spreading analysis

Actin was stained in live imaging experiments with SiR-actin. A threshold was applied to select for actin staining area, with pixel intensity values from approximately 117 (manually adjusted according to staining intensity in each live imaging file) to 255. Actin area in each frame was automatically measured with the Analyze Particles tool in Image J (ImageJ version 2.0.0, MD, USA), selecting for particles above 10 μm^2^.

The number of nuclei or condensates in each frame was measured from Hoechst staining. An automatic threshold was applied and particles from 3 to 3000 μm^2^ were counted in each frame with the Analyze Particles tool. Total actin area was then normalized to nuclei or condensate number in each frame to obtain the area of cell spreading over time.

Actin data from 48 minutes to 15 hours of imaging were fit into a linear model with R Studio (R Studio version 4.0.2, MA, USA).

### 4.8. Measurement of filopodia

Filopodia were measured from phase contrast images at 8 h of live imaging. Images were scouted for any visible filopodia, which were manually measured along their longitudinal axis on Image J (ImageJ version 2.0.0, MD, USA).

### 4.9. Blocking of cell-substrate and cell-cell interactions

For integrin blocking, RGD-Cys-D1 dendrimers were added to the medium to a final concentration of 4×10^−9^ % w/w. We selected this dendrimer concentration because it yields S_18_ substrates. During substrate functionalization, equilibrium is reached between dendrimer concentration in solution and adsorbed dendrimer density; hence, use of the concentration corresponding to the substrates with the lowest density prevents further adsorption mid-assay. Thus, additional dendrimers will not adsorb on the S_90_ substrate but rather attach to free integrin receptors, blocking their interactions with dendrimers on the surface.

To block *N*-Cad and induce gap junction disassembly, ADH1 300 µM or 18βGA 20 µM, respectively, were added to the chondrogenic medium at the start of imaging. We selected those concentrations based on available literature and after observing lighter effects at lower doses but significant cell death at higher doses (ADH1 1000 µM and 18βGA 120 µM).

### 4.10. Statistics

Quantitative data are displayed, showing the mean and standard deviation (SD) or standard error of the mean (SEM) as indicated for each panel. n is the sample size, i.e. the number of tracks analyzed. All experiments were performed at least twice (N = 2), with all data from both computed in the calculations. Data was subjected to a Kolmogorov-Smirnov normality test. For data following a normal distribution, significant differences were judged using the One-way ANOVA with Fisher LSD post-hoc test, or T-test when only two groups were compared. Where data did not pass a normality test, a Kruskal-Wallis’s test with Dunn means comparison was applied. Statistics were performed with OriginPro 8.5 (OriginLab, Madrid, Spain) and GraphPadPrism 8.3 (GraphPad Software, CA, USA).

## Acknowledgements

We thank Dr. Nupur Nagar (University of Vic) for her help with R coding.

## Author contributions

**I. Casanellas:** Conceptualization, Methodology, Investigation, Formal analysis, Visualization, Writing – Original Draft, Writing - Review & Editing. **H. Jiang:** Methodology, Investigation, Formal analysis, Writing - Review & Editing. **C. M. David:** Methodology, Investigation, Formal analysis. **Y. Vida:** Methodology, Resources (design and production of RGD dendrimers), Writing – Review & Editing. **E. Pérez-Inestrosa**: Methodology, Resources (design and production of RGD dendrimers), Writing – Review & Editing, Funding acquisition. **J. Samitier:** Conceptualization, Methodology, Writing – Review & Editing, Supervision, Project administration, Funding acquisition. **A. Lagunas:** Conceptualization, Methodology, Writing – Original Draft, Writing – Review & Editing, Supervision, Project administration, Funding acquisition.

## Data availability

Additional data that support the findings of this study are available from the corresponding author upon request.

